# CancerStop.dev: An Interactive Web Platform Integrating Prognostic Data, Clinical Trials, and Genomic Resources for Patient Empowerment

**DOI:** 10.64898/2026.01.05.697783

**Authors:** Vedanth Ramji, Baladithya Muralitharan, Ganesh Ram, Natarajan Ganesan

## Abstract

Patients facing cancer diagnoses must navigate fragmented information spanning prognosis, clinical research opportunities, and genomics-informed therapy. CancerStop.dev is a React-based web platform that consolidates trusted public resources into a single patient-centered interface to support informed decision-making. The platform’s built-in *ComboReg* module presents interactive relative survival estimates up to 10 years after diagnosis by modeling (combined regression) *age-at-diagnosis* and *stage-at-diagnosis* using publicly available data from the SEER Program. Users can adjust an age slider and view stage-specific curves, enabling individualized and comprehensible visualizations of survivorship trends. A *Clinical Trials* module links directly to ClinicalTrials.gov, providing context-aware queries and free-form keyword filtering (e.g., mutations, investigational agents) to surface ongoing studies. The *Genes & More* module connects to NCBI ClinVar for variant-level insights, facilitating precision medicine exploration when genetic testing results are available. An *Approved Drugs* module routes to the National Cancer Institute resources listing FDA-approved agents relevant to specific cancers, aiding therapy literacy. A curated search tool (*PresciQure*) complements these features by streamlining access to the biomedical literature, package inserts of FDA-approved drugs, and related oncology resources. The platform is non-prescriptive, emphasizes transparency regarding data provenance and limitations, and is positioned to incorporate additional cancer types, demographic stratifications, and multi-omic resources. CancerStop.dev aims to empower patients, caregivers, and clinicians with timely, integrated, and navigable information, thereby strengthening their advocacy and encouraging participation in research and precision care.

**Significance:** CancerStop.dev integrates prognosis, clinical trials, and genomic insights into a single, user-friendly platform, enabling patient and care teams to make faster, better-informed decisions.

## Introduction

Patients and caregivers routinely confront complex treatment choices, survivorship questions, and evolving evidence landscapes following cancer diagnosis. Information relevant to prognosis (e.g., survival expectations), investigational options (e.g., ongoing clinical trials), and genomic findings (e.g., variant-level significance) is commonly distributed across separate systems and formats, creating friction for non-specialists and increasing the cognitive burden at pivotal decision points ^1–3^. Public resources, such as the Surveillance, Epidemiology, and End Results (SEER) Program^4^ for survival statistics, ClinicalTrials.gov^5^ for interventional studies, and NCBI ClinVar^6^ for variant interpretations, are essential, yet they require substantial effort to discover, query, and synthesize^7–9^.

CancerStop.dev^10^ addresses this gap by integrating multiple authoritative sources into a single modern web interface. The platform was initially released as a mobile app (Play Store, 2017) and subsequently rearchitected as a responsive, globally accessible web application built with React. CancerStop.dev^11^ emphasizes clarity of purpose (patient empowerment), transparency of data provenance, and responsible communication (clear disclaimers and non-prescriptive framing) while providing interactive tools to explore relative survival, trials, approved drugs, and variant-level information.

This Resource Report describes the motivation, design, and capabilities of CancerStop.dev, including (i) the *ComboReg* module for age- and stage-informed survivorship curves derived from SEER data, (ii) integrated *ClinicalTrials.gov* querying, (iii) *Genes & More* access to ClinVar for the precision medicine context, (iv) direct routes to *NCI* pages listing FDA-approved agents, and (v) *PresciQure*, a curated search experience for oncology resources. We also outline the usage considerations, limitations, and planned evolution of the resource.

## Implementation

### System Architecture

CancerStop.dev was implemented as a single-page application using React^12,13^, which was designed for dynamic rendering, component reuse, and responsive behavior across device classes [**Figure 1**]. The deployment is hosted on Amazon Web Services (AWS) and configured for scalability, CDN-backed static asset delivery, and robustness under variable traffic. The application employs stateless front-end components and controlled access to external resources through secure HTTPS links and documented APIs, where applicable. Error monitoring and availability checks were used to sustain reliability across modules.

**Figure 1 -.**
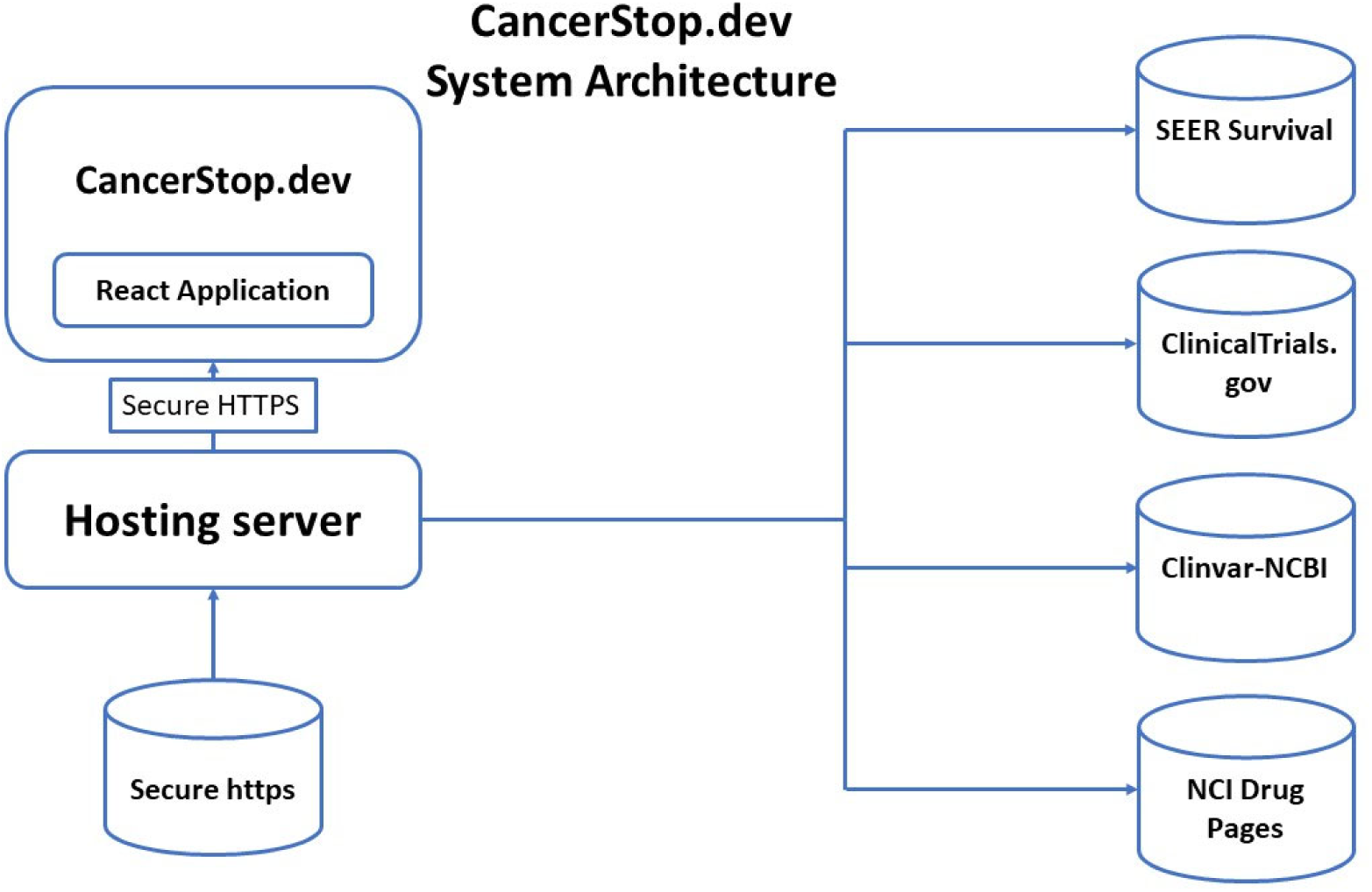
System architecture of CancerStop.dev showing React-based front-end hosted on AWS, with secure HTTPS link-through to SEER, ClinicalTrials.gov, NCBI ClinVar, and NCI drug pages. Stateless components and CDN-backed delivery ensure scalability and responsiveness.

### Design principles

- **Usability and accessibility**: Clear affordances, readable typography, responsive layouts, and descriptive tooltips; alignment with common web accessibility practices. A minimalist approach to the layout was adopted, allowing users to expand upon their chosen modules.
- **Transparency**: Prominent source attribution, contextual help, and domain-specific disclaimers; separation of platform-generated visualizations from original source content.
- **Security and privacy**: No collection of protected health information (PHI) or personally identifiable information (PII); limited, anonymized analytics to monitor performance and usage; HTTPS-only communication.
- **Maintainability**: Modular codebase, versioned deployments, and documentation of data source endpoints.

### Data Sources and Integration Strategy

CancerStop.dev connects users to widely utilized public resources such as SEER^4^, ClinicalTrials.gov^5^, NCBI ClinVar^6^ (variant-level assertions), and NCI drug pages^14^ (FDA-approved agents). Integration is designed to be *link-through* or *contextual query* rather than data warehousing: the platform does not mirror or republish primary datasets; instead, it enables direct access and guided exploration with consistent framing and help content.

- *SEER survivorship*: Source tables present survival by age (18-75) and stage at diagnosis over follow-up intervals (up to 10 years). CancerStop.dev’s ComboReg model interpolates survival estimates for specific ages and presents stage-stratified curves (local/regional/distant/unstaged) using regression/smoothing techniques designed to preserve monotonicity and prevent artifacts near interval boundaries.
- *ClinicalTrials.gov*: Context-aware queries are generated from selected cancer types; the platform exposes a free-form search field for keywords, such as mutations, investigational agents, and locations, with direct navigation to full trial records.
- *NCBI ClinVar*: The ‘Genes & More’ module allows users to input gene names (HGNC/HUGO format) and variant strings (c./g./p. conventions) to reach ClinVar records that summarize assertions and supporting evidence.
- *NCI Approved Drugs*: Cancer-specific lists of FDA-approved agents are presented through direct links to authoritative NCI pages, aiding therapy literacy for patients and caregivers.

### ComboReg Modeling Approach

SEER provides survival statistics in discrete age bands. To offer personalized visualization, CancerStop.dev applies interpolation/regression to estimate survival probabilities for any given age at diagnosis within the covered range. Stage-specific effects are incorporated by applying a second-layer fit to the stage-stratified SEER survival data, yielding four curves that the user can toggle.

#### Stage 1 (age-specific regression)

For each follow-up year, we fitted a smooth regression mapping of the band midpoint (or representative age) to the relative survival S(t). The objective was to interpolate the within-band and between-band gaps to obtain *S*(*t*|age=*a*) for any age within the SEER-supported range. This yielded an age-specific 10-point survival vector, which was rendered as an age-specific survival curve.

##### Constraints and Guardrails

1. Boundedness: 0≤*S*(*t*∣*a*) ≤1
2. Monotonicity in time: *S*(*t*+1∣*a*) ≤*S*(*t*∣*a*)
3. Band continuity: Avoid discontinuities at age-band boundaries via smoothing/regularization.
4. Clinical plausibility: Penalize implausible local oscillations (e.g., survival increases with time).

#### Stage 2 (Stage specific regression from carryover of age-regressed data)

In the second stage, we carried forward age-specific estimates and fit per-stage adjustments to obtain *S* (*t*|*a*, stage). Practically, for each follow-up year *t*, we estimate a stage effect Δ_stage_(*t*∣*a*), such that

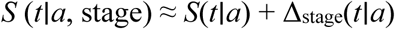

subject to the same boundedness and rank-order guardrails (e.g., distant ≤ regional ≤ local when consistent with aggregates). Alternative parameterizations (multiplicative factors or constrained splines) can be substituted; the UI is agnostic to the internal fit as long as the constraints hold.

The platform displays relative survival up to 10 years post-diagnosis [**Figure 2**] with an interactive age slider and stage-based (where applicable) clear labels referencing SEER’s definitions. The figure also shows a side-by-side comparison of data from the SEER explorer for the selection of cancer type and parameters. The key difference is in the SEER explorer graph; the data are for an age range and stage for a specific selection. In the current implementation, age and stage are combined in a single display, thereby providing a broader picture of relative survival. Implementation prioritizes explainability (help text, FAQs), and includes the guardrails discussed below.

**Figure 2 -.**
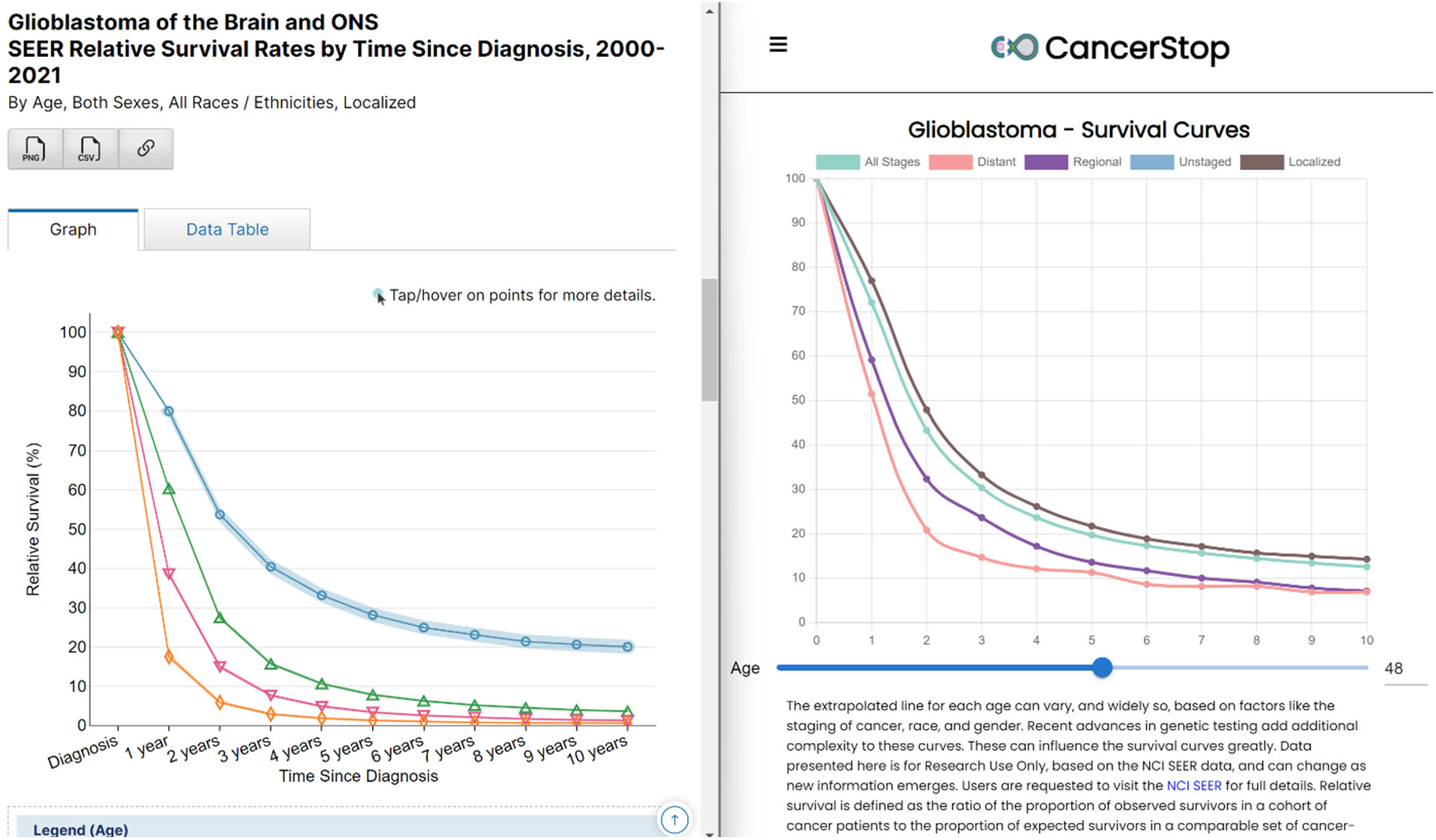
Comparison of SEER Explorer survival plots (age-band and stage-specific) with CancerStop.dev’s ComboReg visualization. CancerStop.dev interpolates survival for a given age and stage using combined regression, presenting all curves in a single interactive display.

### Accessibility and Help Content

The interface is optimized for readability and mobile access; informational popovers and an FAQ page provide instructions, input formats, caveats (e.g., nonprescriptive nature), and links to primary sources. Internationalization and localization are supported at the UI text level, and expanded language coverage is planned for future releases.

### Features and User Experience

The opening page provides a minimalist presentation [**Figure 3**] from where the user can see the cancer modules for which the data are presented. Currently, 15 cancers are presented as modules based on NCI’s SEER Explorer. Sub-classifications for cancers of the Colon and Rectum, Esophagus, Leukemia, and Breast were also included. The list is fully comprehensive and more modules are planned for future versions. However, the menu feature ≡ on the top left of the screen connects the user directly to sites providing relevant information.

**Figure 3 -.**
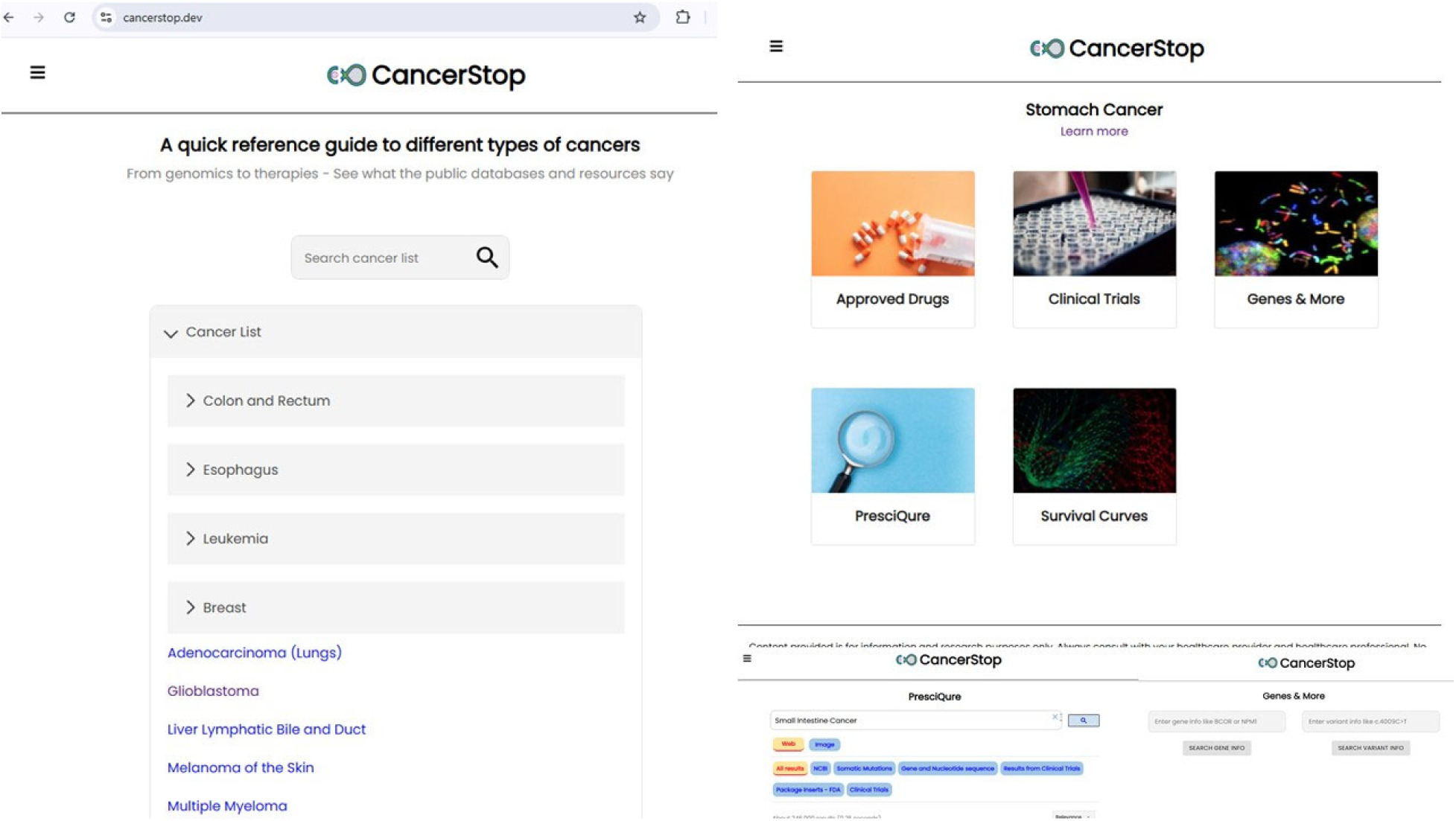
Landing page of CancerStop.dev showing modular tiles for key features: Approved Drugs, Clinical Trials, Genes & More, PresciQure, and Survival Curves. Design emphasizes minimalism and expandability for user-driven navigation.

#### 1) ComboReg Survivorship Data

*Purpose*: To provide ‘relative survival’ estimates tailored to *age-at-diagnosis* and *stage*, supporting comprehension of general survivorship trends without purporting to deliver individual prognosis.

*Interaction*: Users set the diagnosis age with a slider and the plot updates instantly. Survival curves for different tumor stages are presented (local/regional/distance/unstaged). The resultant curves were a result of the combined regression of age and stage-based survival data. Data is modeled from NCI’s SEER Explorer site and hence stage-based presentation is presented where applicable and presented on their site. Tooltips summarize definitions and data provenances.

*Output*: Four stage-specific curves (0–10 years), y-axis labeled “Relative survival,” with clear caveats regarding non-patient-specific interpretation and factors beyond age/stage (e.g., histology, grade, comorbidities, treatment response).

*Rationale*: Convert age-binned SEER data into an approachable visualization for lay stakeholders while preserving the source context and emphasizing limitations.

#### 2) Clinical Trials Integration

*Purpose*: Streamline access to ongoing trials at **ClinicalTrials.gov** is relevant to selected cancers and user-specified terms.

*Interaction*: A prepopulated search targets the selected cancer; a free-form input accepts keywords such as mutations, agents, and trial locations. Advanced filters and full-record views were routed directly to the site via direct navigation.

*Output*: Link-through lists of trial pages, preserving official inclusion/exclusion criteria, contacts, and enrollment information [**Figure 4**].

**Figure 4 -.**
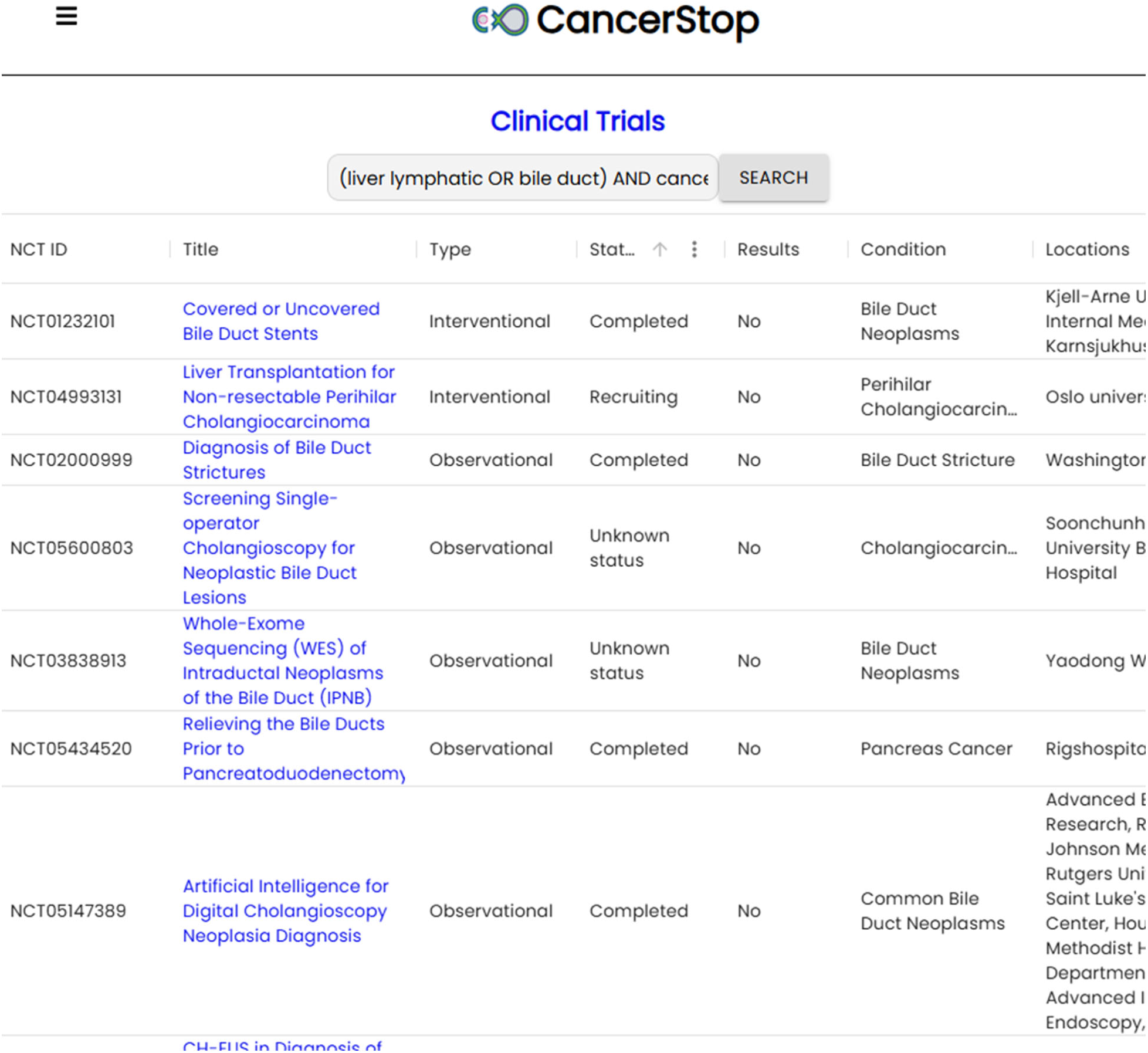
ClinicalTrials.gov integration within CancerStop.dev. Context-aware queries are generated from selected cancer types, with a free-form search field for keywords (e.g., mutations, investigational agents, locations) and direct navigation to official trial records.

*Rationale*: Reduces friction in discovering investigational opportunities and encourages conversations with clinical teams.

#### 3) Genes & More (ClinVar Access)

*Purpose*: Offer variant-level context when users possess genetic testing reports; bridge to ClinVar for assertions and evidence summaries.

*Interaction*: Input gene (HGNC) and variant (c./g./p.) strings, guided examples, and input validation help avoid common formatting errors.

*Output*: Direct navigation to ClinVar records and related links; future expansions intend side-by-side comparisons across resources (e.g., CIViCdb).

*Rationale*: Support precision-medicine literacy without replacing professional interpretations or clinical genetic counseling. Future plans include linking similar resources.

#### 4) Approved Drugs (NCI)

*Purpose*: To connect users to authoritative NCI pages listing FDA-approved agents according to the cancer type.

*Interaction*: Single click from the cancer selection page: The user reviews indications, links to labels, and educational materials.

*Output*: Clear, reliable references for therapy literacy; updates follow NCI editorial cycles.

*Rationale*: Anchors discuss standard-of-care and approved options with a trusted source.

#### 5) PresciQure (Curated Search)

*Purpose*: To provide a curated view of oncology resources (literature, regulatory, trials, and precision oncology) using *Google Custom Search* constrained to vetted domains.

*Interaction*: Tabs refine results (e.g., “*Literature*” for PubMed-aligned content); suggestions in the FAQ invite users to propose additional sources. There are separate tabs that filter results by ‘*Somatic Mutations*’ – connected to CIViCdb^15^, ‘*Package inserts-FDA*’^16^ – connected to publicly available files from the FDA on relevant medications.

*Output*: Focused results that reduce noise; direct links maintain full context and provenance. *Rationale*: Complements platform modules with a lightweight discovery tool aligned with user goals.

### Resource Availability and Access

- **URL**: https://www.cancerstop.dev
- **Access model**: Public, free-to-use, non-prescriptive informational resources, no login required.
- **Code availability**: Source code and deployment scripts are internally hosted on the server; researchers may contact the corresponding author for collaboration inquiries or reproducibility materials (e.g., UI mocks, model specifications).
- **Data handling**: The platform uses link-through and contextual queries and primary data are hosted by their respective providers. CancerStop.dev does not store PHI/PII or republish protected data.

## Results: Usage, Reach, and Illustrative Scenarios

### Adoption and Reach

CancerStop.dev originated as a mobile app in 2017 and transitioned to a web application with steady global visitorship in subsequent years. The platform’s sessions span diverse geographies and device classes, reflecting the need for approachable and integrated views of prognosis and research opportunities. The site and service have also been referenced in the sites and publications^17,18^.

### Illustrative Use Cases

1. Newly diagnosed patients explore age-- and stage-specific relative survival trends, and then navigate to ClinicalTrials.gov to review ongoing studies targeting relevant biomarkers.
2. The caregiver reviews NCI-approved drugs for a cancer subtype and uses *PresciQure* to surface educational materials for shared decision-making.
3. Clinician or navigator references *ClinVar* entries for a reported variant to guide discussions about evidence strength and potential trial eligibility, emphasizing that the final interpretation requires professional judgment.

### User Safeguards

Across modules, CancerStop.dev integrates *help text*, *FAQs*, and *disclaimers* emphasizing (i) informational use only, (ii) need for clinician consultation, (iii) heterogeneity across patients (e.g., histology, grade, prior therapy), and (iv) non-substitutability for medical advice.

### Validation and Quality Assurance

#### Visual and Numerical Checks

ComboReg outputs are evaluated to maintain (i) *monotone non-increasing* survival with increasing follow-up, (ii) curve smoothness that respects the SEER interval structure, (iii) *boundedness* in [0,1], and (iv) *stage rank-order coherence* consistent with published aggregates. Visual inspections and synthetic edge-case tests (e.g., ages near band boundaries) were used to detect artifacts.

#### Data Provenance and Updates

Because CancerStop.dev primarily links to authoritative resources, content updates reflect upstream changes (e.g., newly posted trials and revised drug pages). SEER-based visualizations are revalidated when upstream tables or definitions change (e.g., revised survival cohorts or stratifications). The FAQ points users to source documentation to contextualize terms like “*Relative survival*.”

## Discussion

### Contribution

CancerStop.dev provides a *unified, patient-centered entry point* for core public oncology resources, lowering barriers to exploring survival expectations, investigational options, and variant-level contexts. By coupling interactive visualization (ComboReg) with direct navigation (ClinicalTrials.gov, NCI drugs, ClinVar) and a curated search layer (PresciQure), the platform promotes timely access to information and shared decision-making.

### Distinguishing Characteristics (contextual)

CancerStop.dev differs from resource-specific portals in that it orchestrates multiple sources within a consistent UI and framing. It avoids data duplication, emphasizing provenance, link-through transparency, and non-prescriptive messaging. This orientation is suited to patients and caregivers seeking orientation rather than expert-level analytic workflows.

### Limitations

- *Population-level statistics*: Relative survival curves derived from SEER represent cohorts and may not apply to individual patients; many covariates (e.g., grade and therapy) are not modeled in the current visualization.
- *Variant interpretation*: ClinVar records vary in assertion strength and may contain conflicts, and professional genetic counseling continues to remain essential.
- *Trials navigation*: Eligibility depends on detailed criteria; trial availability is dynamic and location-dependent.
- *Scope and curation*: PresciQure’s curated domains are evolving, and comprehensive coverage will continue to be evaluated and included in the list.
- *Localization*: Although the UI is responsive, expanded language support is ongoing.

### Future Directions

Planned enhancements include:

- *Expanded stratifications*: Incorporation of additional covariates for survival (e.g., with and without intervention from a specific treatment, sex, histology, race/ethnicity) where source data permit, presented with clear caveats.
- *Precision integrations*: Side-by-side views of variant evidence across multiple databases (e.g., CIViCdb), with educational context on actionability.
- *Guided pathways*: Wizard-like flows help users formulate trial queries and understand inclusion and exclusion criteria.
- *Open collaboration*: opportunities for external contributions to curation lists, UX testing, and localization.

### Ethics, Disclaimers, and Governance

CancerStop.dev is an information resource. They do not provide medical advice, diagnoses, or treatment recommendations. Users are encouraged to consult healthcare professionals regarding decisions related to diagnosis, therapy, and trial participation. The platform prominently displays disclaimers and source attributions. Governance emphasizes privacy (no PHI/PII collection), security (HTTPS only), and responsible communication (clear caveats and help content).

### Author Contributions

Conceptualization: NG, VR, BM, GR; Methodology (ComboReg modeling): NG, VR; Software (React implementation): VR; Curation (PresciQure): NG, VR, BM; Resources and data integration: NG, VR, BM; Writing—original draft: NG; Writing—review and editing: all authors; Supervision: NG.

### Data and Code Availability

CancerStop.dev provides link-through access to SEER, ClinicalTrials.gov, NCI drug pages, and NCBI ClinVar. The platform does not redistribute the upstream datasets. The model specifications and UI artifacts may be shared upon reasonable request from the corresponding author.

### Ethics Statement

This study did not involve human participants, animals, or identifiable personal data.

